# CRISPR activation screening identifies VGLL3 and GATA2 as transcriptional regulators of PD-L1

**DOI:** 10.1101/2021.08.18.456771

**Authors:** Ruud H. Wijdeven, Birol Cabukusta, Xueer Qiu, Daniel M. Borras, Yun Liang, Jacques Neefjes

## Abstract

The PD-L1/2 – PD-1 immune checkpoint is essential for the proper induction of peripheral tolerance and limits autoimmunity, whereas tumor cells exploit their expression to promote immune evasion. Many different cell types express PD-L1/2, either constitutively or upon stimulation, but the factors driving this expression are often not defined. Here, using genome-wide CRISPR-activation screening, we identified three factors that upregulate PD-L1 expression; GATA2, MBD6, and VGLL3. GATA2 and VGLL3 act as transcriptional regulators and their expression induced PD-L1 in many different cell types. Conversely, loss of VGLL3 impaired IFNγ-induced PD-L1/2 expression in keratinocytes. Mechanistically, by performing a second screen to identify proteins acting together with VGLL3, we found that VGLL3 forms a complex with TEAD1 and RUNX1/3 to drive expression of PD-L1/2. Collectively, our work identified a new transcriptional network controlling PD-L1/2 expression and suggests that VGLL3, in addition to its known role in the expression of pro-inflammatory genes, can balance inflammation by upregulating the anti-inflammatory factors PD-L1 and PD-L2.

## Introduction

The engagement of inhibitory checkpoint receptor PD-1 on CD8+ T-cells by its ligands PD-L1 and PD-L2 impairs T-cell activation and effectively dampens the immune response (1, 2). Many different cell types express PD-L1 and to a lesser extent PD-L2, including tumour cells, immune cells, endothelial cells and epithelial cells (3–5). Tumour cells express PD-L1 to promote immune evasion and inhibition of the PD-1/PD-L1 checkpoint is now widely used as immunotherapy against many different tumour types (6, 7). On the other hand, immune cells and non-immune cells, such as keratinocytes, express PD-L1 to induce peripheral tolerance and the absence of PD-L1 on these cells leads to excessive immune activation and autoimmunity (2, 8–11).

Many cell types express PD-L1 constitutively at low levels and this is boosted by pro-inflammatory stimuli, such as Toll-like receptor activators and interferon-γ (IFNγ). Similar to many other immune genes, the PD-L1 promoter contains binding sites for AP-1, NF⍰B, Myc, STAT and IRF1, the latter being responsible for IFNγ-induced expression activation of PD-L1 (12–14). Tumour cell-specific expression of PD-L1 can be mediated via these transcription elements, but also via copy number alterations, mRNA stabilization or 3’ UTR shortening (15–18). Alternative mechanisms inducing PD-L1 expression include activation via YAP/TAZ, HIF1α or ATF3 factors (19–22). However, these factors fail to explain the differential expression in all tumour cells. Furthermore, the mechanisms driving PD-L1 expression in specific subtypes of immune cells and non-immune cells remain unknown. Recent genome-wide knock-out screens for cell surface expression of PD-L1 have identified CMTM4 and CMTM6 as regulators of PD-L1 protein stability (23, 24), but no new transcriptional regulators.

To identify novel regulators of PD-L1 surface expression, we conducted a genome-wide CRISPR activation screen. This screen yielded two transcriptional regulators of PD-L1: GATA2 and VGLL3. GATA2 utilizes its DNA binding capacity to drive PD-L1 and PD-L2 transcription, whereas VGLL3 forms a complex with TEAD1 and RUNX1/RUNX3 to mediate PD-L1 expression. GATA2 and VGLL3 function in cooperation with IFNγ and we show that VGLL3 is required for proper induction of PD-L1 and PD-L2 by IFNγ in keratinocytes. Together, our work identifies new regulatory proteins involved in the expression of the PD-L1 and PD-L2 immune checkpoint ligands.

## Results

### A CRISPR activation screen identifies GATA2, MBD6, and VGLL3 as regulators of PD-L1 expression

The immune checkpoint inhibitor PD-L1 is expressed in many tissues at varying levels, suggesting that several mechanisms are in play controlling its expression. To identify novel regulators promoting PD-L1 cell surface expression, MelJuSo cells were transduced with a genome-wide CRISPR/Cas9-mediated gene activation library targeting 23,000 genes of the human genome (25, 26) (Fig. 1A). This human cutaneous melanoma cell line expresses low levels of PD-L1, which provides a window for detecting increased expression. Following transduction with the activation library, cells with increased PD-L1 surface expression were FACS-sorted twice and analyzed for gRNA enrichment compared to the input (Fig. 1A). Hits were considered based on two criteria: two or more gRNAs per gene showed 4-fold enrichment in the sorted population in two independent sorts, and this enrichment had to be statistically significant according to RSA analysis (Fig. 1B and S1A). Based on these criteria, several hits were identified, with the top one being CD274, the gene encoding PD-L1. To validate these results, cell lines stably expressing individual gRNAs were generated. Analysis of PD-L1 surface expression in these cells revealed that guides activating GATA2, MBD6 and VGLL3 upregulated PD-L1 expression (Fig. 1C). Meanwhile, these guides did not affect expression of another cell surface marker, MHC class I. Indeed, these gRNAs upregulated the expression of their respective target genes (Fig. S1B); identifying the transcription factor GATA2, transcriptional co-activator VGLL3, and polycomb-binding protein MBD6 as regulators of PD-L1 cell surface expression in the melanoma cell line MelJuSo.

**Figure 1:**
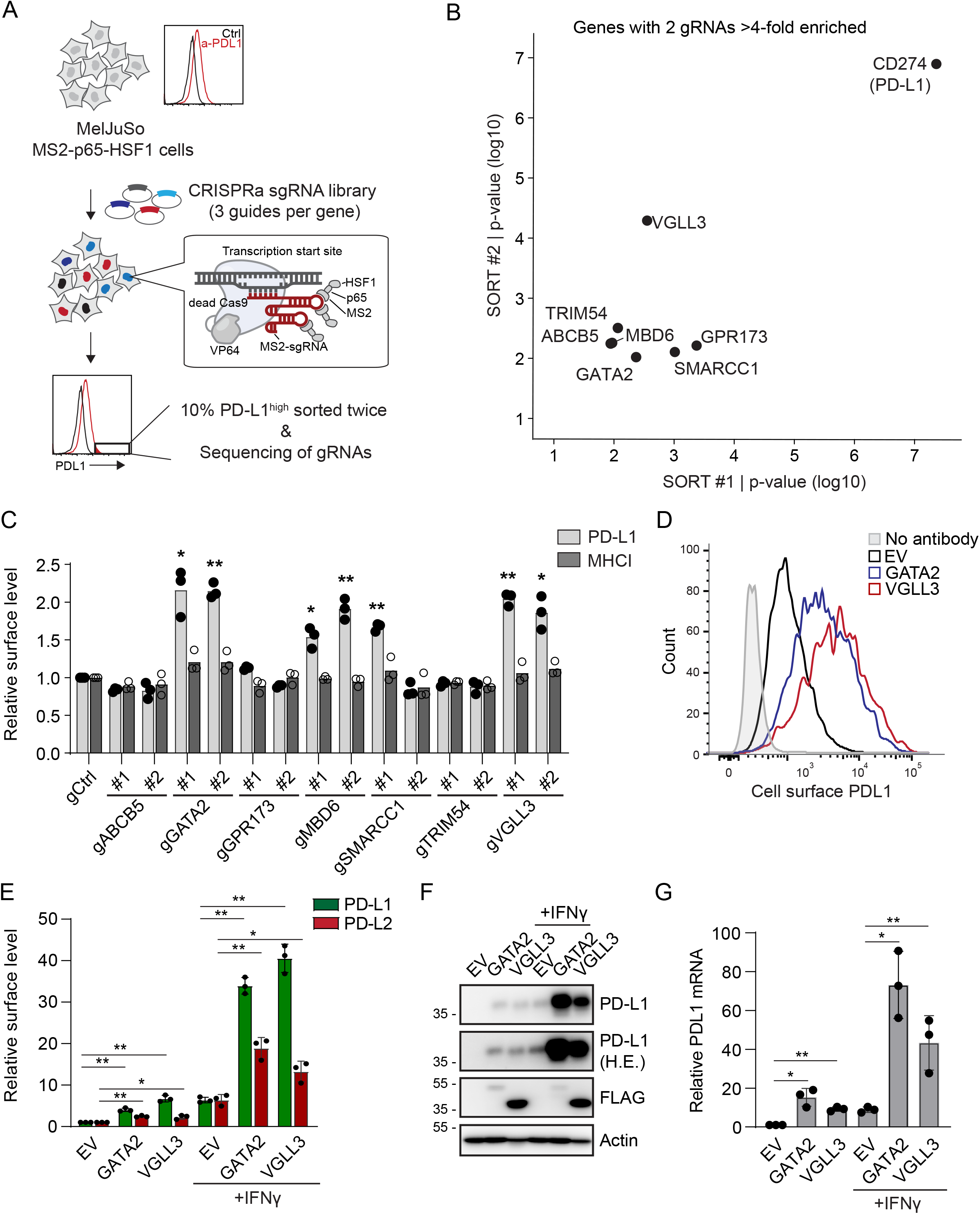
CRISPRa screen identifies novel regulators of PD-L1 expression. (A) Schematic set-up of the screen. MelJuSo melanoma cells stably expressing MS2-p65-HSF1 were transduced with a pooled gRNA library containing dCAS9 and FACS-sorted for cells displaying high levels of PD-L1. (B) Genes for which at least two different gRNAs were significantly enriched (> 4-fold) in the sorted population versus control population in both replicate sorts. Plotted are p-values based on RSA analysis. (C) MelJuSo MPH cells stably expressing the SAM vector with or without the indicated activation gRNAs were analyzed for cell surface expression of PD-L1 and MHC class I (HLA-ABC). Data represent three independent experiments (+SD), statistical significance was determined by a paired student’s T-test (*, P <0.05, **, P <0.01). (D) MelJuSo cells stably expressing FLAG (EV), GATA2-FLAG or FLAG-VGLL3 were analyzed for cell surface expression of PD-L1 using flow cytometry. (E) MelJuSo cells as in (D) were either or not stimulated with IFNγ for 48h and cell surface expression of PD-L1 and PD-L2 was measured using flow cytometry. (F) MelJuSo cells as in (D) were either or not stimulated with IFNγ for 24h and expression of the indicated proteins was determined by Western blot. (G) MelJuSo cells as in (D) were treated with IFNγ for 24h when indicated and mRNA levels of the indicated genes were analyzed using qRT-PCR and normalized to GAPDH. All data represent three independent experiments (+SD), statistical significance was determined by a paired student’s T-test (*, P <0.05, **, P <0.01).

### GATA2 and VGLL3 regulate PD-L1 transcription

Due to their defined functions as transcriptional regulators, we followed-up on GATA2 and VGLL3. To confirm gRNA specificity and eliminate the possibility of off-target effects, PD-L1 expression was assessed after over-expressing GATA2 and VGLL3 using cDNA expression plasmids. Both FLAG-tagged GATA2 and VGLL3 robustly upregulated expression of PD-L1 at the cell surface, protein, and mRNA level (Fig. 1D-G). Interestingly, IFNγ stimulation of GATA2 and VGLL3-overexpressing cells further boosted their PD-L1 expression, suggesting that GATA2 and VGLL3 cooperate with IFNγ in promoting PD-L1 expression (Fig. 1E-G). GATA2 and VGLL3 also upregulated the expression of the PD-L1 homologue PD-L2 (Fig. 1E). Increased PD-L1 mRNA could be the consequence of mRNA stabilization or increased transcription. To test for the first, cells were cultured with transcriptional inhibitor actinomycin D, showing that GATA2 or VGLL3 did not affect the half-life of PD-L1 mRNA (Fig. S1C). Overall, our results demonstrate that GATA2 and VGLL3 promote transcription of PD-L1 and PD-L2, a finding which for GATA2 was recently shown by others in glioblastoma cells as well (27).

Mutating the DNA binding site for GATA2 abolished its ability to upregulate PD-L1, arguing it acts as a direct transcription factor (Fig. S1D). When transducing different cell lines, GATA2 was found to upregulate PD-L1 in many (but not all) cell types, including all non-tumorigenic cell lines tested (Figure 3A). GATA2 is expressed by multiple cell types that express PD-L1 constitutively, such as different immune cell types and endothelial cells, suggesting a more general role for GATA2 in PD-L1 expression.

**Figure 2:**
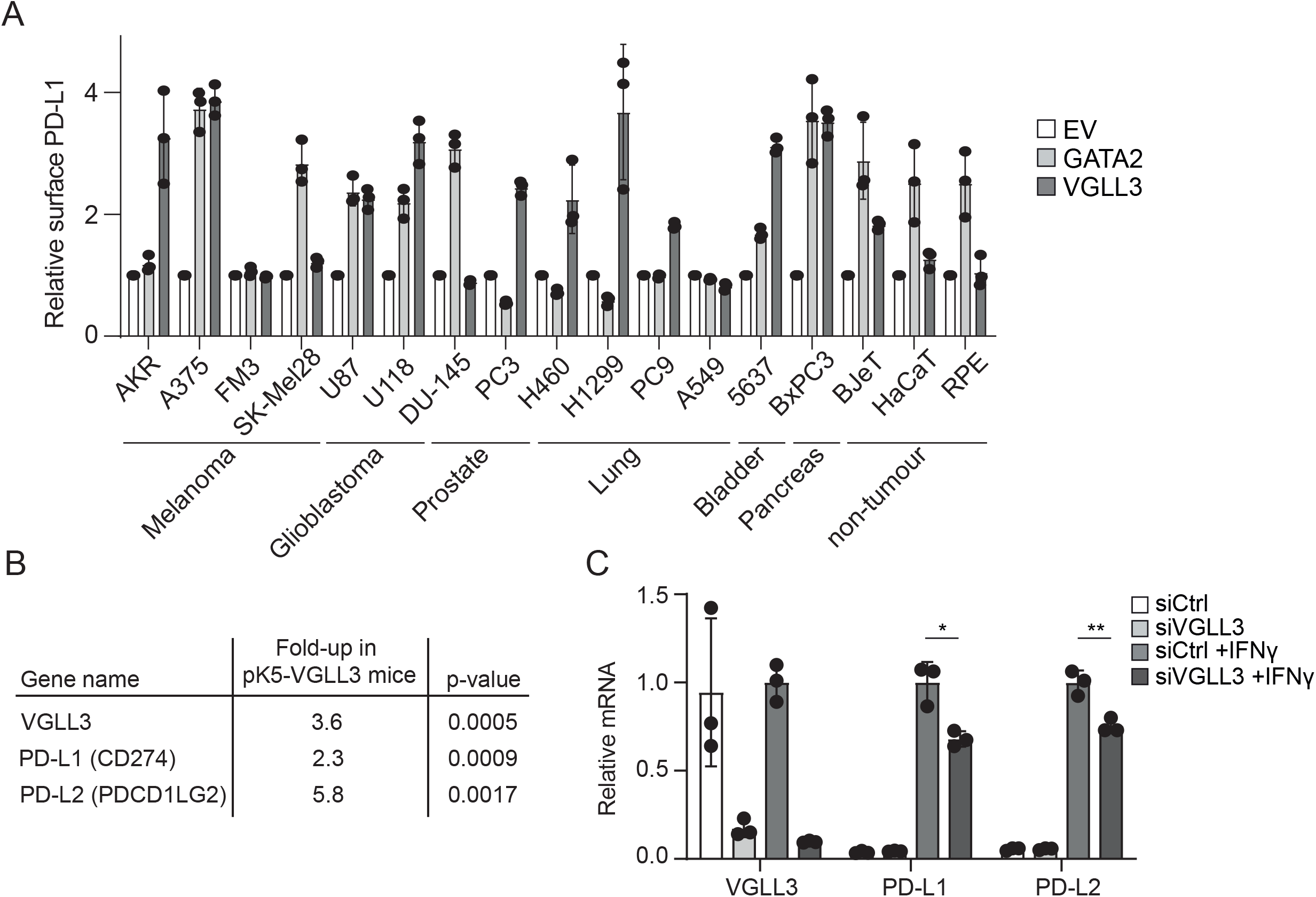
VGLL3 regulates IFNy-induced PD-L1 expression in keratinocytes. (A) The different cell lines were transduced with FLAG, GATA2-FLAG, or FLAG-VGLL3 cDNA constructs. Four days later, cells were stained for PD-L1 and analyzed by flow cytometry. GFP+ cells were gated and PD-L1 expression was normalized to the Flag-transduced cells. Cells corresponded to different tissues. (B) Upregulation of VGLL3, PD-L1, and PD-L2 transcripts in the skin of mice overexpressing VGLL3 from a transcriptome data set (29). (C) Keratinocytes were transfected with siCtrl or siVGLL3 and mRNA for the indicated genes was analyzed from cells either or not treated with IFNy for 48h. Shown are data from three independent experiments (+SD), statistical significance determined by a paired student’s T-test (*, P <0.05, **, P <0.01).

**Figure 3:**
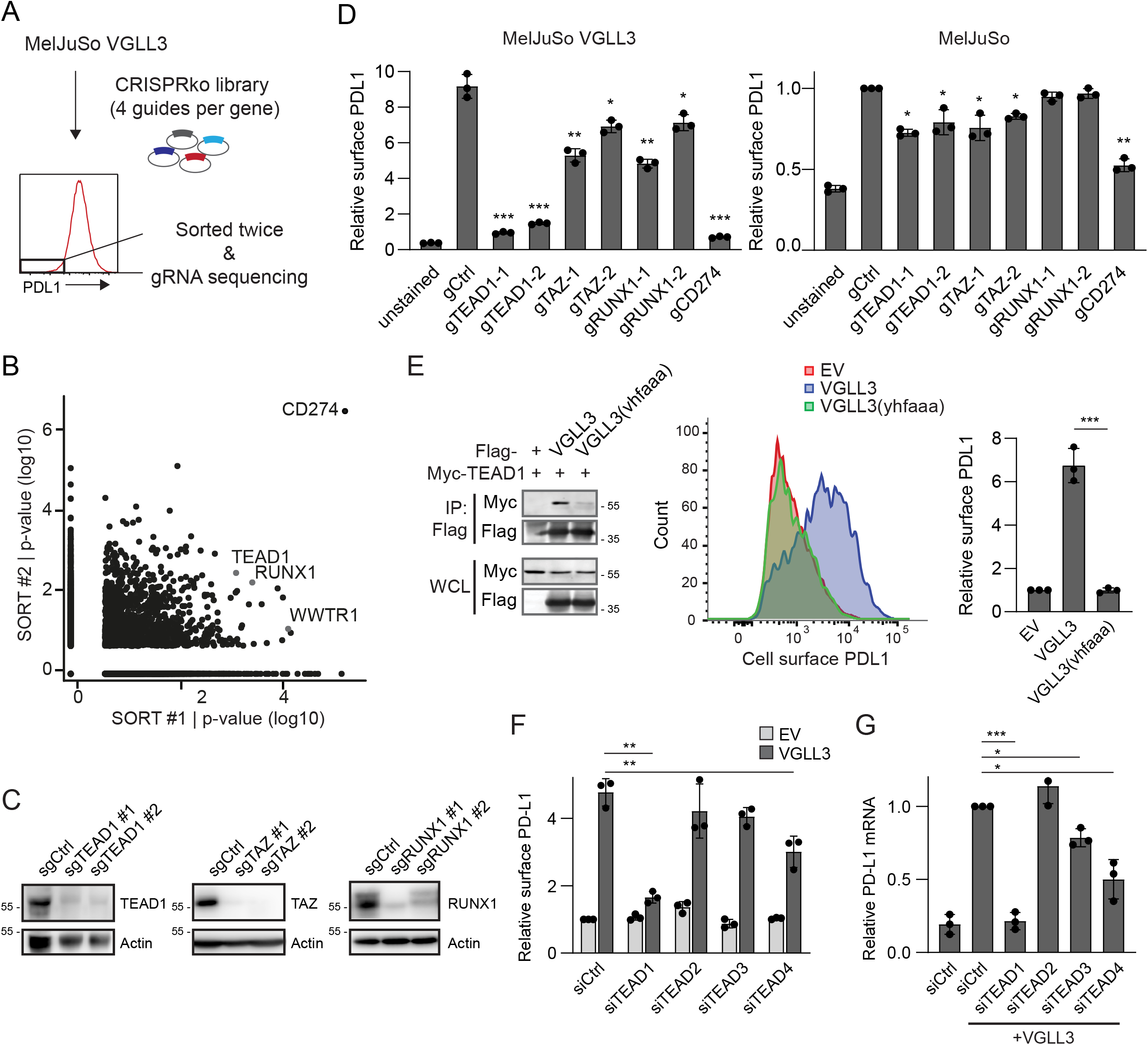
VGLL3 cooperates with TEAD1 to drive PD-L1 expression. (A) Schematic set-up of the screen. MelJuSo cells stably expressing FLAG-VGLL3 were transduced with the Brunello CRISPR knockout library and FACS-sorted twice for cells displaying low PD-L1 surface levels. (B) Results of the RSA analysis of the inserts from the biological duplicates, with three candidates indicated with grey dots. (C) Western blot validation of the knockout efficiency of the pooled MelJuSo VGLL3 knockout cells transduced with the indicated gRNAs. (D) MelJuSo FLAG-VGLL3 or FLAG expressing cells were transduced with the indicated gRNAs and pooled knockout lines were analyzed for surface PD-L1 expression using flow cytometry. (E) Left: Myc or Myc-TEAD1 were isolated from HEK293T cells using Myc-TRAP beads and associated FLAG-VGLL3 or FLAG-VGLL3(vhfaaa) was detected by Western blot. Right: MelJuSo cells transduced with the indicated expression constructs were analyzed for expression of PD-L1 using flow cytometry. (F) MelJuSo cells stably expressing FLAG or FLAG-VGLL3 were transfected with the indicated siRNAs and three days later analyzed for PD-L1 expression using flow cytometry. (G) As in (F) but three days after siRNA transfection. mRNA was isolated and the expression of PD-L1 transcript was analyzed by qRT-PCR and normalized to GAPDH mRNA. All data represent three independent experiments (+SD), statistical significance determined by a paired student’s T-test (*, P <0.05, **, P <0.01, ***, P <0.001).

### VGLL3 regulates IFNγ-induced PD-L1 and PD-L2 expression in keratinocytes

VGLL3 is a transcriptional activator that has recently been shown to drive expression of several pro-inflammatory genes and is reported to be involved in the induction of autoimmunity in women (28–30). Our data suggest that VGLL3 also promotes the expression of the anti-inflammatory genes PD-L1 and PD-L2. Similar to GATA2, many, but not all, cell types upregulated PD-L1 in response to VGLL3 expression (Fig. 2A). Interestingly, HaCaT keratinocytes were somewhat sensitive to VGLL3, and the previous studies were also performed in keratinocytes, arguing VGLL3 can regulate PD-L1 and PD-L2 expression in keratinocytes. In agreement, mice overexpressing VGLL3 specifically in keratinocytes (under the Keratin-5 promoter) displayed higher levels of PD-L1 and PD-L2 mRNA (29) (Fig. 3B). To validate that VGLL3 controls PD-L1/2 expression in these cells, primary keratinocytes were depleted for VGLL3 using RNAi. While VGLL3 depletion did not affect the basal levels of PD-L1/2 expression in these cells, IFNγ-induced expression of PD-L1/2 was hampered when VGLL3 was depleted (Fig. 2C). Overall, these results show that VGLL3 is an activator of IFNγ-induced expression of PD-L1 and PD-L2 in keratinocytes.

### VGLL3 controls PD-L1 transcription via TEAD1

While VGLL3 upregulated PD-L1 in many cell types, the fact that not all cell types responded suggests that VGLL3 alone is not sufficient to promote PD-L1 expression, i.e. it requires additional (transcriptional) factors. To identify partners cooperating with VGLL3 in driving PD-L1 transcription, we performed a genome-wide CRISPR knockout screen in the MelJuSo FLAG-VGLL3 expressing cells, sorting for cells with low PD-L1 expression (Fig. 4A).

**Figure 4:**
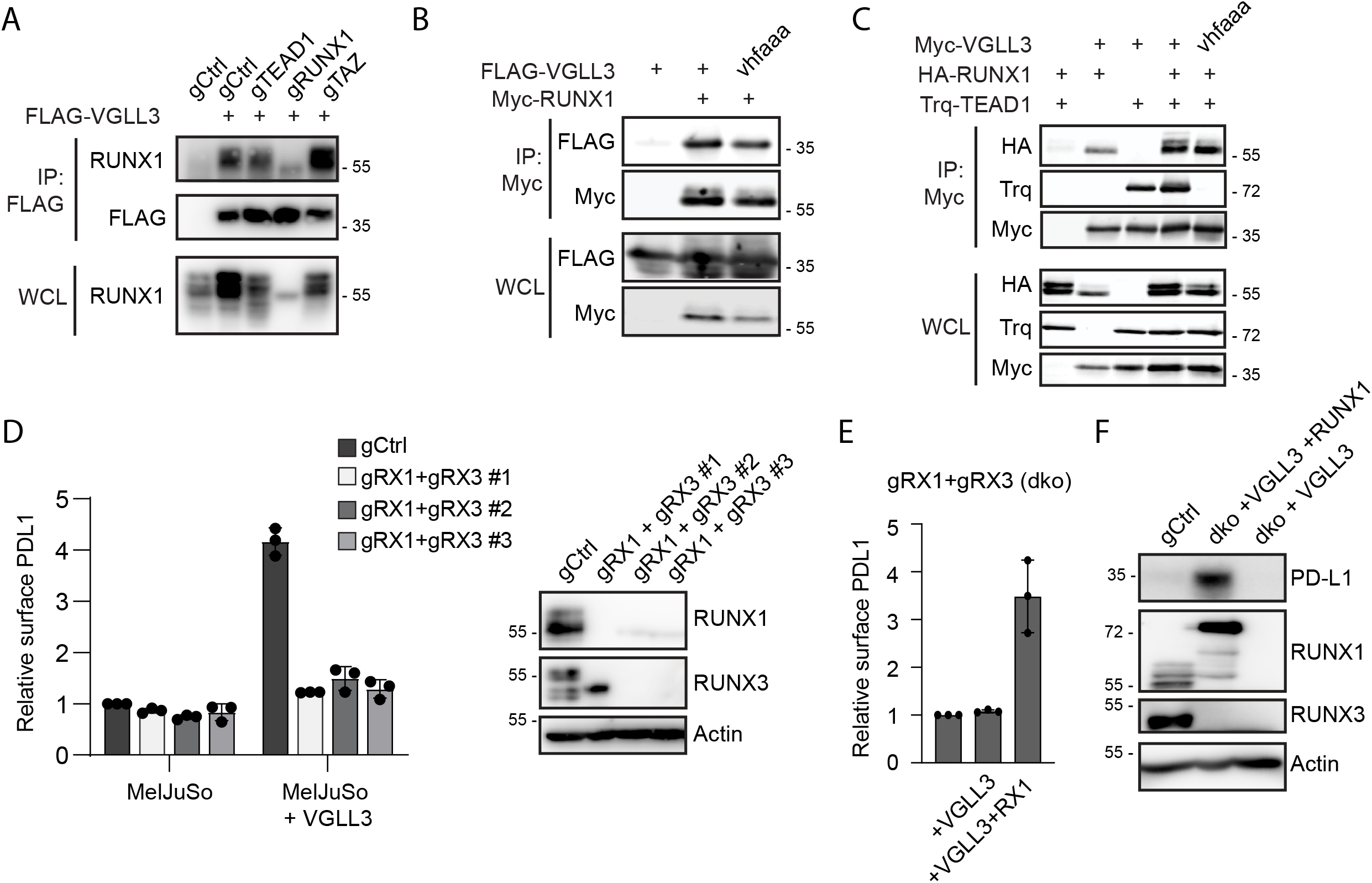
VGLL3 interacts with RUNX1/3 to drive PD-L1 expression. (A) MelJuSo cells with the indicated genetic background were lysed, and FLAG or FLAG-VGLL3 was immunoprecipitated. Associated RUNX1 was detected by Western blot. (B) Myc or Myc-RUNX1 were isolated from HEK293T cell lysates using Myc-TRAP beads and associated Flag-VGLL3 or Flag-VGLL3-vhfaaa mutant was detected by Western blot. (C) Myc or Myc-VGLL3 were isolated from HEK293T cell lysates using Myc-TRAP beads and associated HA-RUNX1 or Turquoise(Trq)-TEAD1 was detected by Western blot. (D) MelJuSo cells or single cell knockout clones of MelJuSo cells transduced with RUNX1 and RUNX3 targeting sgRNAs were either or not transduced with Flag-VGLL3 and PD-L1 surface expression was analyzed by flow cytometry. (E) MelJuSo cells depleted for RUNX1 and RUNX3 were transfected with Flag-VGLL3 and Trq-RUNX1 when indicated and cell surface expression of PD-L1 was analyzed by flow cytometry. (F) MelJuSo cells either or not depleted for RUNX1 and RUNX3 (double knockout, dko) were transduced with FLAG-VGLL3 and Trq-RUNX1 when indicated and the indicated proteins were detected by Western blot.

gRNA enrichment analysis of the sorted cells identified CD274 as the top hit, substantiating the screen (Fig. 3B). In addition to CD274, gRNAs targeting the nuclear factors RUNX1, TAZ (WWTR1), and TEAD1 were significantly enriched (Fig. S2A). It is reported that TAZ regulates PD-L1 expression in many tumor cells, in conjunction with the TEAD transcription factors (21, 22). Moreover, VGLL3 interacts with TEAD transcription factors via its TDU domain to act as a transcriptional co-activator (31). This suggests VGLL3 interacts with TEAD1 to drive PD-L1 expression, in a manner similar to TAZ. Indeed, gRNA-mediated knockout of TEAD1 in FLAG-VGLL3 cells almost completely abrogated VGLL3-mediated PD-L1 expression (Fig. 3C-D). Meanwhile, the effect of TEAD1 knockout on constitutive PD-L1 expression in MelJuSo cells was less prominent (Fig. 3D, right panel). On the other hand, the effect of TAZ knockout on PD-L1 surface expression was similar with or without VGLL3 overexpression (Fig. 4D). This argues that TAZ is involved in PD-L1 expression in MelJuSo cells and acts independently from VGLL3. Lastly, we found that the knockout of RUNX1 did not affect the existing PD-L1 expression but reduced the VGLL3-mediated expression by about 50%.

To validate that the interaction between VGLL3 and TEAD1 is critical to drive PD-L1 expression, we generated a TDU domain mutant that in other VGLLs abolishes TEAD-binding (V166A, H167A, F170A). This mutation indeed prevented VGLL3 from interacting with TEAD1 and failed to upregulate PD-L1 (Fig. 3E). In contrast, mutating a Histidine-stretch that is required for VGLL3 to control transcription in association with the ETS1 transcription factors did not show any effect on VGLL3-controlled PD-L1 expression (Fig. S2B) (32). In line, RNAi-mediated depletion of TEAD1 and to a lesser extent TEAD4 decreased VGLL3-dependent upregulation of PD-L1, even though the expression of TEAD4 was much higher in these cells (Fig. 4F, 4G and S2C). Thus, VGLL3 interacts with TEAD1 to drive PD-L1 expression, in a manner similar to the oncogenic factor TAZ. VGLL1 did not upregulate PD-L1, so the ability to upregulate PD-L1 is not conserved in all VGLLs (Fig. S2D).

### VGLL3 interacts with RUNX1 and RUNX3 to drive PD-L1 expression

Our knockout screen also identified RUNX1 as a candidate involved in VGLL3-mediated PD-L1 expression. To test whether VGLL3 interacts with RUNX1, VGLL3 was immunoprecipitated from lysates. RUNX1 co-precipitated with VGLL3, and the absence of TAZ and TEAD1 expression did not affect this interaction (Fig. 4A), suggesting that VGLL3 interacts with RUNX1 independently of TEAD1. Indeed, TDU-mutant VGLL3 interacted as strongly as wild-type VGLL3 with RUNX1 (Fig. 4B and 4C). Thus, TEAD1, RUNX1 and VGLL3 form a tripartite complex to drive PD-L1 expression. However, while TEAD1 was essential for VGLL3-driven PD-L1 expression, RUNX1 depletion only affected this partly. RUNX1 is homologous to and shares redundancy with RUNX3 (33), so we tested whether co-depletion of RUNX1 and RUNX3 would render cells insensitive to VGLL3. Double-knockout clones were generated using RUNX1 sgRNA #1 in combination with three different RUNX3 CRISPR constructs. Each of these clones failed to upregulate PD-L1 expression after VGLL3 transduction (Fig. 4D). Sensitivity of these cells for VGLL3 was restored by ectopic expression of RUNX1 (Fig. 4E and 4F), indicating that RUNX1 and RUNX3 act redundantly in conjunction with VGLL3. Collectively, we propose that VGLL3 forms a transcriptional complex with TEAD1 and RUNX1/RUNX3 in control of PD-L1 expression.

## Discussion

The PD-L1/2 – PD-1 immune checkpoint is essential for the induction of peripheral tolerance and limiting autoimmunity, whereas tumour cells exploit it to promote immune evasion. Given the wide variety of cell types expressing PD-L1, and the major differences in expression levels, it is likely that many factors are involved in the control of PD-L1/2 expression. Several factors that independently drive PD-L1 transcription have been identified, but research has mostly focused on the pathways relevant in tumor cells. Here, we used an unbiased CRISPR-activation approach to identify novel factors regulating PD-L1. By doing so, factors usually not expressed in the model cell line are included, although it is limited to factors that can drive expression independently, or in the genetic context of the chosen cell line. In addition to the recently published transcription factor GATA2, we identified MBD6 and VGLL3 as novel regulators of PD-L1 expression. Whereas MBD6, a polycomb-interacting protein, was not further studied, VGLL3 was shown to complex with TEAD1 and RUNX1/3 to regulate PD-L1 transcription.

While cytokine-induced PD-L1 expression and tumor cell expression of PD-L1 are studied in detail, it is unknown which factors drive constitutive expression in different immune cell subsets and other cell types such as endothelial cells and keratinocytes. Our data in combination with a study by Fu Y et al. (27) show that GATA2 facilitates PD-L1 expression. GATA2 is predominantly expressed in various immune cell types, as well as endothelial cells, which both have significant PD-L1 expression. Furthermore, GATA2 is highly expressed in hematopoietic stem cells, like PD-L1/2, and their expression correlates during the different phases of stem cell development (34). It will be interesting to test whether GATA2 is indeed responsible for PD-L1/2 expression during hematopoiesis and what the physiological function is for this expression.

We also identified VGLL3 as a novel regulator of both PD-L1 and PD-L2 expression. VGLL3 is a transcriptional co-activator that can induce transcription via the TEAD transcription factors, as well as ETS1 (31, 32). We define a tripartite transcription complex consisting of VGLL3, TEAD1 and RUNX1/3 in the control of PD-L1 expression. The TEAD-transcription factors contain four members and although in our system TEAD1 appears to be dominant, it could be that in different cell types the other TEAD-factors act together with VGLL3. Similarly, RUNX1 and RUNX3 are very homologous and act redundantly to afford PD-L1 induction. The prerequisite for RUNX1/3 sets VGLL3 apart from YAP/TAZ, which also drive PD-L1 expression via the TEAD-family (21, 22, 35, 36). However, RUNX1 and RUNX3 also interact with these factors, so VGLL3 and TAZ potentially act mechanistically similarly to drive PD-L1 expression. VGLL1 on the other hand did not induce PD-L1 expression, and VGLL4 has been reported to induce PD-L1 independently from its interaction with the TEADs (37). This suggests that within the VGLL-family VGLL3 has specifically evolved to regulate PD-L1.

VGLL3 is a transcriptional activator that displays a female-dominant expression and is linked to several autoimmune diseases that are more prevalent in women (28). It regulates the expression of many autoimmune-associated pro-inflammatory genes, and overexpression in keratinocytes leads to the development of a lupus-like rash and systemic autoimmunity (28–30). Our data demonstrate that complementary to promoting expression of pro-inflammatory genes, VGLL3 induces expression of the anti-inflammatory factors PD-L1 and PD-L2. Using the same models as before, mice expressing VGLL3 in keratinocytes were found upregulate PD-L1/2, and keratinocytes were shown to require VGLL3 for full induction of PD-L1/PD-L2 by IFNγ. Thus, VGLL3 promotes expression of genes from both sides of the pro- and anti-inflammatory equation. This resembles IFNγ, which also promotes pro-inflammatory cytokines such as CXCL9 and CXCL10, as well as anti-inflammatory factors like PD-L1 and PD-L2. Based on these data, we propose that VGLL3 serves as an additional factor to control PD-L1/L2 expression under conditions of inflammation. This then occurs at sites where VGLL3 is expressed. In mice, the expression of VGLL3 is highest in trophoblasts, especially extravillous trophoblasts and syncytiotrophoblasts (Figure S3) (38, 39), which form the outer layer and protrusions of the placenta. These are also among the highest PD-L1 and PD-L2 expressing cells, and secrete pro-inflammatory factors to recruit and shape an immune cell repertoire that maximally protects the fetus (40). It is tempting to speculate that these tissues utilize VGLL3 to create such a well-balanced but active immune environment.

In summary, using a genome-wide CRISPR activation screen in combination with a dedicated genome-wide CRISPR knock-out screen, we identified novel regulators of PD-L1 expression, including GATA2 and MBD6. VGLL3, a female-biased transcriptional activator linked to autoimmune disease, in complex with TEAD1 and RUNX1/3 was shown to regulate transcription of PD-L1 and PD-L2. VGLL3 is required for full induction of PD-L1/2 expression by IFNγ in keratinoctyes and thus not only regulates expression of pro-inflammatory genes, but also expression of the anti-inflammatory immune checkpoint molecules PD-L1 and PD-L2.

## Materials and methods

### Cell culture

MelJuSo cells were cultured in IMDM supplemented with 8% fetal calf serum and cell line authentication was performed by Eurofins Genomics (19-ZE-000487). HEK 293T cells were obtained from the ATCC (CRL-3216) and cultured in DMEM supplemented with 8% fetal calf serum. AKR, A375, FM3, SK-MEL-28, U87, U118, A549, BJeT, HaCaT, and RPE were cultured in DMEM supplemented with 8% serum; DU145, PC3, H460, H1299, PC9, 5637, and BxPC3 were cultured in RPMI supplemented with 8% serum. Normal primary human keratinocytes were purchased from Lonza (Morristown, NJ, USA) and cultured in Lonza KGM-gold keratinocyte growth medium according to manufacturer’s instructions. One day before cytokine stimulation, cells were switched to Lonza KBM medium to remove growth factors. Keratinocytes were used at passage 1 or 2.

### Transfections, transductions and antibodies

For the generation of viral particles, HEK 293T cells were transfected using polyethyleneimine (Polyscience Inc.) with packaging plasmids pRSVrev, pHCMV-G VSV-G and pMDLg/pRRE in combination with the lentiviral construct. Virus was harvested, filtered and target cells were transduced in the presence of 8μg/ml polybrene (Millipore). For co-immunoprecipitation experiments, HEK 293T were also transfected using polyethyleneimine.

For siRNA mediated depletion, cells were reverse transfected with DharmaFECT transfection reagent #1 and 50 nM siRNA (catalog numbers: siCtrl: D00120613-20, siTEAD1: M-012603-01-0005, siTEAD2: M-012611-00-0005, siTEAD3: M-012604-01-0005, siTEAD4: M-019570-03-0005) according to the manufacturer’s protocol. Briefly, siRNAs and DharmaFECT were mixed and incubated for 20 minutes in a culture well, after which cells were added and left to adhere. Cells were analyzed three days after siRNA transfections. For gene depletion in keratinocytes, cells were electroporated using Lonza 4D-nucleofector following manufacturer’s recommendations.

Antibodies used in this study are as following: Flow cytometry: PE anti-human CD274 (BioLegend), PE anti-human PD-L2 (BioLegend), FITC anti-human HLA-ABC (Biolegend). Western blot: mouse anti-PD-L1 (Cell Signaling, #29122), mouse anti-Actin (Sigma, A5441), mouse anti-FLAG (Sigma, M2), rabbit anti-TEAD1 (ProSci 22-472), mouse anti-TAZ (Santa Cruz, sc-518026), mouse anti-RUNX1 (Santa Cruz, sc-365644), mouse anti-RUNX3 (Santa Cruz, sc-101553), mouse anti-HA (Covance 16B12), rabbit anti-GFP and mouse anti-Myc (9E10), both were described in (41).

### CRISPR-activation and knockout screen

For CRISPR activation, the human CRISPR 2-plasmid activation pooled library (SAM) was a gift from Feng Zhang (Addgene #1000000078) and used for CRISPR activation screening. Stable MPH-expressing MelJuSo cells were generated and 150 million MelJuSo MPH cells were infected at an MOI of 0.3 with the SAM plasmid. The next day, cells were selected with hygromycin (200μg/ml) and blasticidin (2.5μg/ml) and after five days, two batches of cells were stained for PD-L1 and the top 10% of PD-L1 expressing cells were sorted out using the FACS. Cells were grown out for another six days and PD-L1^high^ cells were sorted again for both replicates. After the second sort, cells were grown out to reach 10 million and gDNAs were isolated and amplified using the established protocol (25). gRNAs were sequenced using the Illumina Highseq 2500 and inserts were mapped to the reference. Statistical analysis was done using RSA analysis, enrichment >4 was considered a hit (42).

For knockout screening, the human CRISPR Brunello genome-wide knockout library was a gift from David Root and John Doench (Addgene #73178). MelJuSo cells stably expressing FLAG-VGLL3 were generated and two batches of 100 million cells were infected at an MOI of 0.3. Transduced cells were selected using puromycin (1μg/ml) and after five days, cells were stained for PD-L1 and the lowest 5% of GFP-positive PD-L1^low^ expressing cells were sorted. Cells were grown out and sorted again using the same gating strategy as for the first sort. After this sort, cells were grown out to reach 10 million, the genomic DNA was isolated and gDNAs were amplified using the established protocol (25). gRNAs were sequenced using the Illumina NovaSeq6000 and inserts were mapped to the reference.

### Hit validation and subcloning

For hit validation, two individual guides per gene were cloned into the LentiSAM vector (a gift from Feng Zhang, Addgene plasmid #75112) and MelJuSo MPH cells were stably transduced with these guides or with the empty LentiSAM as a control. Guide sequences used were as follows: GATA2-1: 5’- GGACTCCGGGACTGACCCGC-3’, GATA2-2 5’-GTCCGCAATTCCCGAACCGC-3’, MBD6-1 5’- GGCTCCTGCGGCGGCGGCTG-3’, MBD6-2 5’-TGCGCAGTGCCTTCTGGGAA-3’, VGLL3-1 5’- GGGGAACGCCGGGGAGAAGG-3’, VGLL3-2 5’-TGCCTCCCCAGGCTCTGACG-3’, ABCB5-1 5’- -3’, ABCB5-2 5’- -3’, GPR173-1: 5’- -3’, GPR173-2 5’- -3’, SMARCC1-1: 5’- GATGCAGCCTACGGAGGATGG-3’, SMARCC1-2: 5’- GTCAAAAACCGGCGCTCACTA-3’, TRIM54-1 5’- -3’, TRIM54-2 5’- -3’.

For validation of the knockout screen, two individual guides per gene were cloned into the LentiCRISPR v2 vector (a gift from Feng Zhang, Addgene plasmid #52961) and the indicated MelJuSo cells were transduced and selected using puromycin. Guide sequences were as follows: CD274: 5’- GGTTCCCAAGGACCTATATG-3’, TAZ-1: 5’- ATCCGAAGCCTAGCTCGTGG-3’, TAZ-2: 5’- -ACGCGGGCGACGAGTGCGAG-3’, TEAD1-1: 5’- ACATGGTGGATAGATAGCCA-3’, TEAD1-2: 5’- GGCCGGGAATGATTCAAACA-3’, RUNX1-1: 5’- CACTTCGACCGACAAACCTG-3’, RUNX1-2: 5’- TAGATGATCAGACCAAGCCC-3’, RUNX3-1: 5’- CCCCAGGATGCATTATCCCG-3’, RUNX3-2: 5’- CACTGCGGCCCACGAAGCGA-3’, RUNX3-3: 5’- CCGTGCCGTACCTTGGATTG-3’.

### Expression plasmids

Expression constructs for GATA2-FLAG (NM_001145661.2), FLAG-VGLL3 (NM_016206.4), FLAG-VGLL1, and GATA2Δ14 (NM_001145662) and VGLL3 mutants were made using Gateway cloning of the indicated genes into the pLenti-CAG-gate-FLAG-IRES-GFP vector, which was a gift from William Kaelin (Addgene plasmid #107398). For GATA2 (C373R), VGLL3 VHFAAA (V166A, H167A, F170A), VGLL3 ∆HIS (∆242-248) the mutagenesis primers as depicted in table 1 were used. Turquoise (Trq)-RUNX1, Trq-TEAD1, Myc-VGLL3 and HA-RUNX1 were generated by subcloning these genes into the Trq-C1, 2Myc-C1 or 2HA-C1 vectors.

**Table 1.**
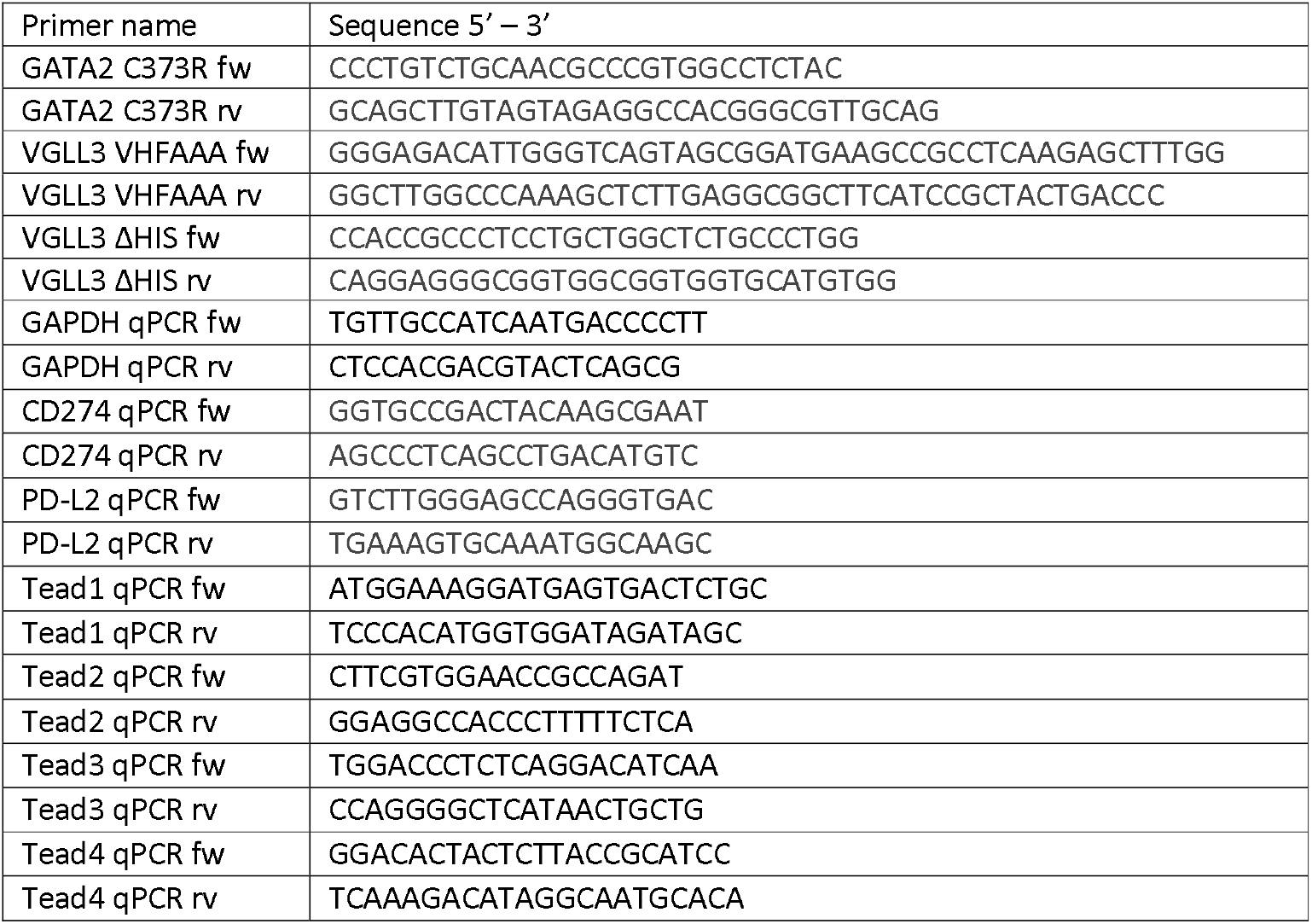
Primer sequences

### Flow cytometry

Cells were trypsinized and stained for the indicated antibodies in 2% fetal calf serum PBS. After washing, cells were analyzed on the BD LSRII and data was analyzed using FlowJo software.

### cDNA synthesis and qPCR

RNA isolation (Bioline) and cDNA synthesis (Roche) were performed according to the manufacturer’s instructions. SYBR green (Bioline) signal was detected on the BIO-RAD analyzer and normalized to GAPDH using the Pfaffl formula. Primers used for detection of signals can be found in Table 1. Actinomycin D was purchased from Santa Cruz Biotechnology.

### Co-immunoprecipitation and Western blotting

For co-immunoprecipitation experiments, cells were lysed in lysis buffer (0.5% NP-40, 5% glycerol, 150mM NaCl, 50mM Tris-HCl pH8.0, 5mM MgCl_2_ supplemented with complete EDTA-free Protease Inhibitor Cocktail (Roche)) and cleared by centrifugation. Lysates were incubated with Myc-Trap beads (Chromotek) or protein G-Sepharose 4 FF resin pre-loaded with the indicated antibodies. Following incubations, beads were washed extensively with lysis buffer before addition of SDS-sample buffer (2% SDS, 10% glycerol, 5% β-mercaptoethanol, 60mM Tris-HCl pH 6.8 and 0.01% bromophenol blue).

For whole cell lysate analyses, cells were lysed directly in SDS-sample buffer. Samples were boiled before loading and proteins were separated by SDS-PAGE and transferred to Western Blot filters. Blocking of the filter and antibody incubations were done in PBS supplemented with 0.1 (v/v)% Tween20 and 5% (w/v) milk powder. Blots were imaged using the Odyssey Imaging System (LI-COR) or Amersham Imager.

## Acknowledgements

We thank the LUMC Flow Cytometry Facility for support and cell sorting, and members of the Neefjes group for critical discussions. This work was supported by the Institute of Chemical Immunology (ICI), an NWO Gravitation project funded by the Ministry of Education, Culture and Science of the government of the Netherlands, a Spinoza Premium and a European Research Council (ERC) advanced grant awarded to J.N.

## Notes

### Competing Interest Statement

The authors have declared no competing interest.

