## Supplemental figures for "CRISPR activation screening identifies VGLL3 and GATA2 as transcriptional regulators of PD-L1"

### Supplemental data

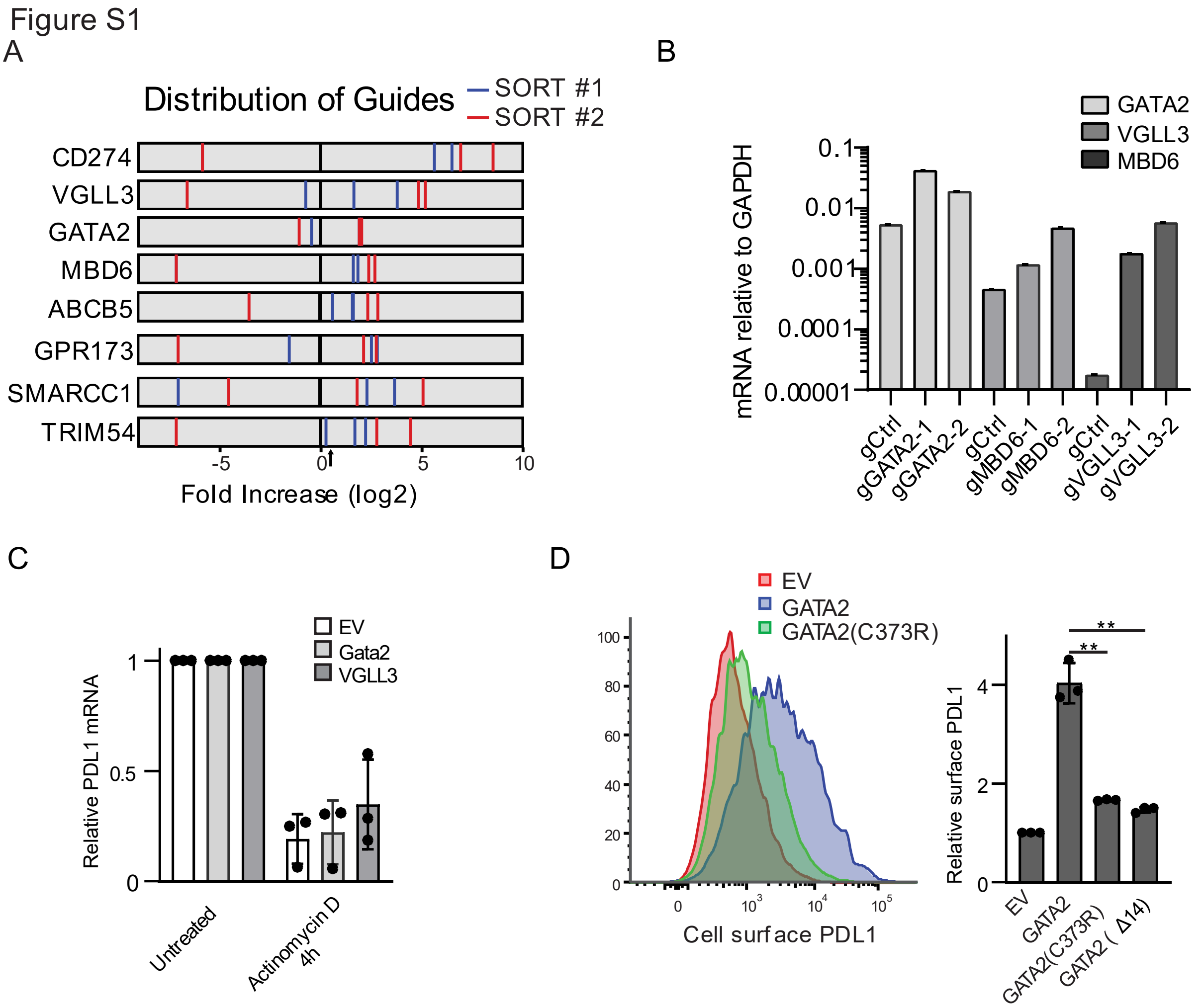


**Figure S1: CRISPRa screen identifies novel regulators of PD-L1 expression, related to Figure 1.** (A) gRNA enrichment for hits with at least two guides per sort enriched >4-fold. Arrow depicts 4-fold threshold and red and blue stripes represent enrichment of individual gRNAs compared to the non-sorted population., (B) RNA was isolated from MelJuSo MPH cells stably expressing the indicated gRNAs and expression of the respective target genes was analyzed using qRT-PCR. Expression was normalized to GAPDH. (C) MelJuSo cells stably expressing FLAG (EV), GATA2-FLAG or FLAG-VGLL3 were either or not treated with 1µM Actinomycin D for 4h to block transcription. mRNA levels of PD-L1 were measured at indicated timepoints using qRT-PCR and normalized to GAPDH and the respective non-treated control. (D) MelJuSo cells transfected with the indicated cDNA expression constructs were analyzed for expression of PD-L1 using flow cytometry. Data from (C) and (D) represent three independent experiments (+SD), statistical significance was determined by a paired student's T-test (**, P <0.01).

Figure S2

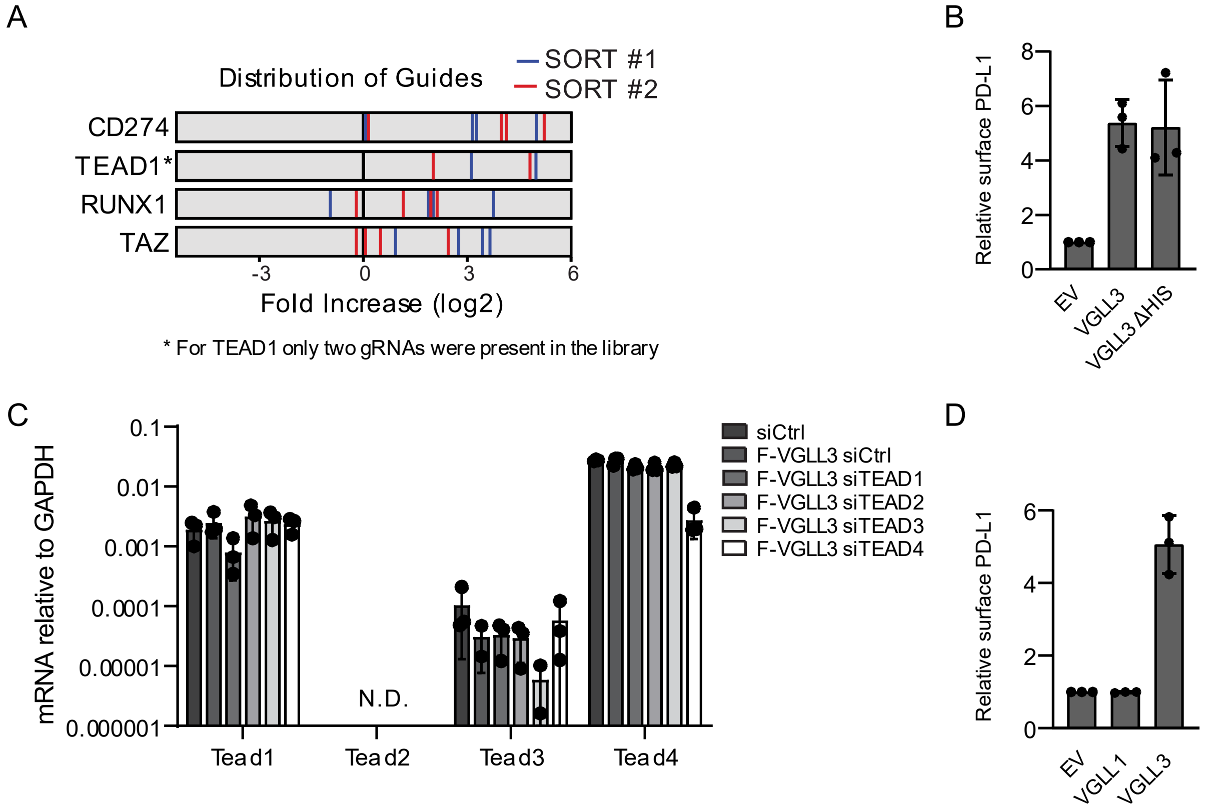


**Figure S2: VGLL3 cooperates with TEAD1 to drive PD-L1 expression, related to Figure 4.** (A) gRNA enrichment for indicated hits from the knockout screen. Red and blue stripes represent enrichment of individual gRNAs compared to the non-sorted population. (B) MelJuSo cells transduced with the indicated expression constructs were analyzed for expression of PD-L1 by flow cytometry. (C) MelJuSo cells stably expressing FLAG or FLAG-VGLL3 were transfected with the indicated siRNAs. Three days later, mRNA was isolated and the expression of TEAD transcripts was analyzed by qRT-PCR and normalized to GAPDH mRNA levels. (D) MelJuSo cells were transduced with FLAG, FLAG-VGLL1 or FLAG-VGLL3 cDNA expression constructs and PD-L1 expression was measured by flow cytometry. All data represent three independent experiments (+SD).

Figure S3 From Human Protein Atlas (proteinatlas.org)

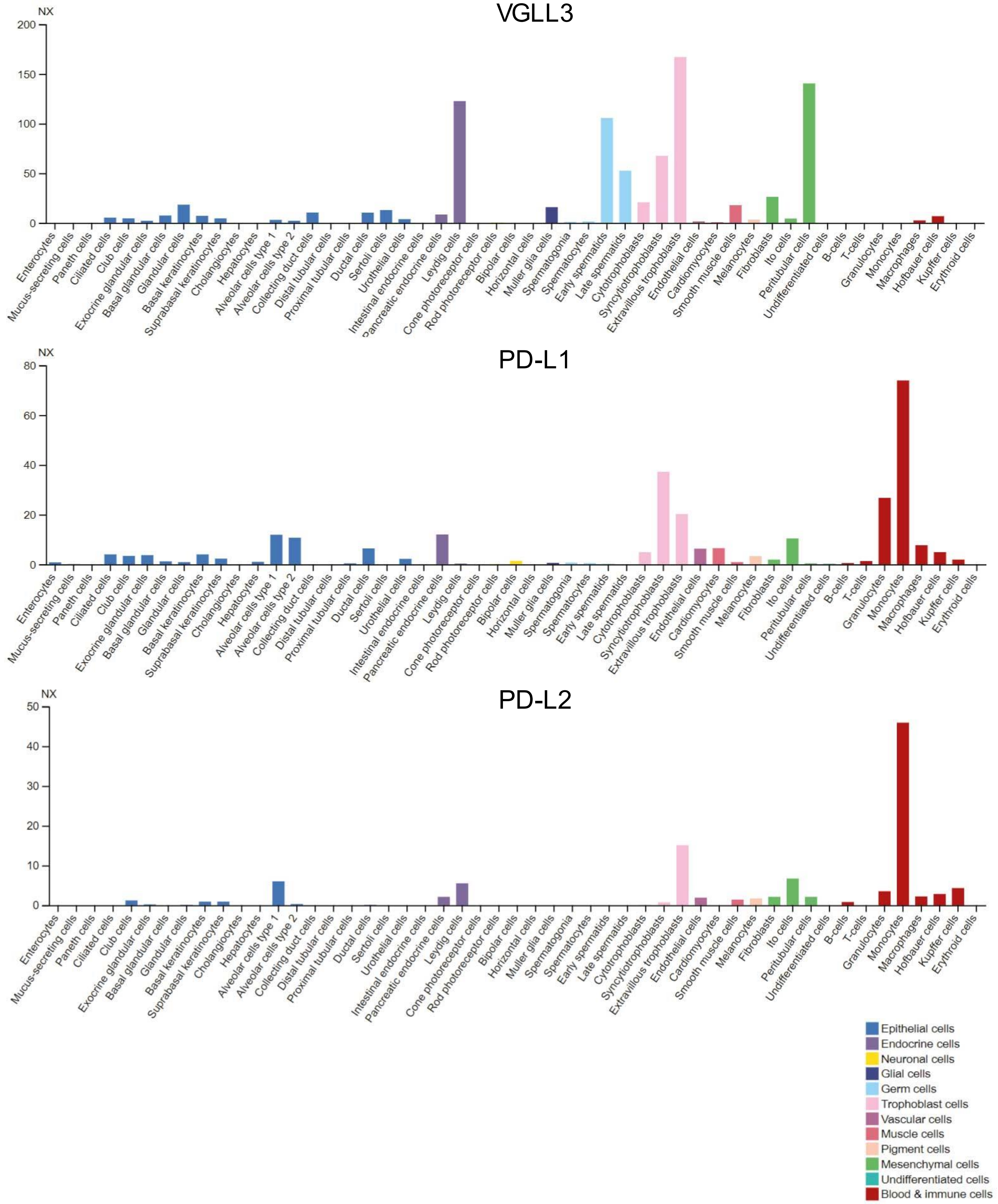


**Figure S3: VGLL3 and PD-L1/2 are highly expressed in trophoblasts.** mRNA expression data from different cell types for VGLL3, PD-L1 and PD-L2 were extracted from the human protein atlas database. In pink the three types of trophoblasts. Other cell types displaying high levels of VGLL3 transcripts are all located in the male reproductive organ (Leydig cells, peritubular cells and spermatids).
